# Additional analytical support for a new method to compute the likelihood of diversification models

**DOI:** 10.1101/693176

**Authors:** Giovanni Laudanno, Bart Haegeman, Rampal S. Etienne

**Affiliations:** Groningen Institute for Evolutionary Life Sciences, Box 11103, 9700 CC, Groningen, The Netherlands; Centre for Biodiversity Theory and Modelling, Theoretical and Experimental Ecology Station, CNRS and Paul Sabatier University, Moulis, France

**Keywords:** Diversification, Birth-death process

## Abstract

Molecular phylogenies have been increasingly recognized as an important source of information on species diversification. For many models of macro-evolution, analytical likelihood formulas have been derived to infer macro-evolutionary parameters from phylogenies. A few years ago, a general framework to numerically compute such likelihood formulas was proposed, which accommodates models that allow speciation and/or extinction rates to depend on diversity. This framework calculates the likelihood as the probability of the diversification process being consistent with the phylogeny from the root to the tips. However, while some readers found the framework presented in Etienne et al. (2012) convincing, others still questioned it (personal communication), despite *numerical* evidence that for special cases the framework yields the same (i.e. within double precision) numerical value for the likelihood as analytical formulas do that were independently derived for these special cases. Here we prove *analytically* that the likelihoods calculated in the new framework are correct for all special cases with known analytical likelihood formula. Our results thus add substantial mathematical support for the overall coherence of the general framework.

## 1 Introduction

One of the major challenges in the field of macro-evolution is understanding the mechanisms underlying patterns of diversity and diversification. A very fruitful approach has been to model macro-evolution as a birth-death process which reduces the problem to the specification of macroevolutionary events (i.e. speciation and extinction). However, providing likelihood expressions for these models given empirical data on speciation and extinction events is quite challenging, for the following reason. While such a likelihood is very easy to derive when full information is available for all events, typically the data involves phylogenetic trees constructed with molecular data collected from extant species alone. Hence, no extinction events and speciation events leading to extinct species are recorded in such phylogenetic trees. For a variety of models this problem can be overcome by considering a reconstructed process, whereby the phylogeny of extant species can be regarded as a pure-birth process with time-dependent speciation rate [9]. But this approach is not generally valid.

Thus, the methods employed to derive likelihood expressions are usually applicable to a limited set of models. They do not apply to models that assume that speciation and extinction rates depend on the number of species in the system. Hence, potential feedback of diversity itself on diversification rates, due to interspecific competition or niche filling, is completely ignored. The first to incorporate such feedbacks were Rabosky & Lovette [10], who made rates dependent on the number of species present at every given moment in time, analogously to logistic growth models used in population biology. However, their model had to assume that there is no extinction for mathematical tractability, which stands in stark contrast to the empirical data: the fossil record provides us with many examples of extinct species.

Etienne et al. [3] presented a framework to compute the likelihood of phylogenetic branching times under a diversity-dependent diversification process that explicitly accounts for the influence of species that are not in the phylogeny, because they have become extinct. Some of our colleagues have doubts that this framework contains a formal argument that the solution of the set of ordinary differential equations that together constitute the framework gives the likelihood of the model for a given phylogenetic tree. Instead, only numerical evidence for a small set of parameter combinations has been provided that the method yields, in the appropriate limit, the known likelihood for the standard diversity-independent (i.e. using constant-rates) birth-death model. This likelihood was first provided by Nee et al. [9], using a breaking-the-tree approach. Later, Lambert & Stadler [8] used coalescent point process theory to provide an approach to obtain likelihood formulas for a wider set of models. These models did not include diversity-dependence. For example, Lambert et al. [4, 7] applied their framework to the protracted birth-death model, which is a generalization of the diversity-independent model where speciation is no longer an instantaneous event [5]. For this model they provided an explicit likelihood expression.

Here we provide an analytical proof that the likelihood of Etienne et al. [3] reduces to the likelihood of Lambert et al. [7] – and hence to that of Nee et al. [9] – for the case of diversity-independent diversification.

The extant species belonging to a clade are often not all available for sequencing, because some species are difficult to obtain tissue from (either because they are difficult to find/catch, or because they are endangered, or because they have recently become extinct due to anthropogenic rather than natural causes) or because it is difficult to extract their DNA. This means that our data consists of a phylogenetic tree of an incomplete sample of species, and thus of an incomplete set of speciation events, even for those that lead to the species that we observe today. Models have been developed to deal with this incomplete sampling. One model assumes that we know exactly how many extant species are missing from the phylogeny. This is the case for well-described taxonomic groups, such as birds, where we have a good idea of the species that are evolutionarily related, but we are simply missing some data points for the reasons mentioned above. This sampling model is called *n*-sampling [7]. Another model assumes that we do not know exactly how many species are missing from the phylogeny, but rather that we have an idea of the probability *ρ* of a species being sampled. This sampling scheme is called *ρ*-sampling [7], but is also referred to as *f*-sampling [9]. The framework of Etienne et al. [3] assumes *n*-sampling, but in this paper we show that it can also be extended to incorporate *ρ*-sampling.

In the next section we summarize the framework of Etienne et al. [3] and we provide the likelihood formula analytically derived by Lambert et al. [7] for the special case of diversity-independent but time-dependent diversification with *n*-sampling. Then we proceed by showing that the probability generating functions of these two likelihoods are identical. We end with a discussion where we point out how the framework of Etienne et al. [3] can be extended to include *ρ*-sampling and how it relates to the likelihood formula of Rabosky & Lovette [10] for the diversity-dependent birthdeath model without extinction.

## 2 The diversity-dependent diversification model

Diversification models are birth-death processes in which “birth” and “death” correspond to speciation and extinction events, respectively. In the simplest case, the per-species speciation rate *λ* and the per-species extinction rate *μ* are constants. Here we consider diversification models in which the speciation and extinction rates depend on the number of species *n* present at time *t*, i.e., diversity-dependent, which we denote by *λ*_*n*_ and *μ*_*n*_. We also allow the speciation and extinction rates to depend on time *t*, i.e., *λ*_*n*_(*t*) and *μ*_*n*_(*t*), although the latter dependence is often not explicit in our notation.

We assume that the diversification process starts at time *t_c_* from a crown, i.e., from two ancestor species. Assuming that at a later time *t* > *t*_*c*_ the process has *n* species, the transition probabilities in the infinitesimal time interval [*t*, *t* + *dt*] are

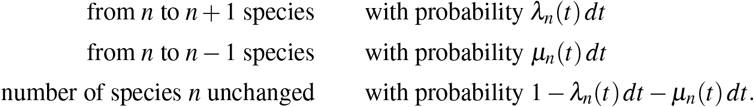

The diversification process runs until the present time *t*_*p*_.

We denote by *P*_*n*_(*t*) the probability that the process has *n* species at time *t*. This probability satisfies the following ordinary differential equation (ODE, called master equation or forward Kolmogorov equation [1]),

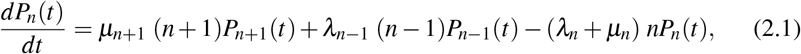

where we omit in the notation the time dependence of the speciation and extinction rates.

### Sampling models

At the present time *t*_*p*_ a subset of the *n* extant species are observed and sampled. This sampling process can been modelled in two different ways (see introduction). The first model assumes that a fixed number of species is unsampled, which corresponds to the *n*-sampling scheme of Ref. [7]. That is, the number of extant species at *t*_*p*_ that are not sampled, a number we denote by *m*_*p*_, is part of the data. The second model assumes that each extant species at the present time is sampled with a given probability, which has been called *f*-sampling [9] or *ρ*-sampling [7]. In this case the number of unsampled species *m*_*p*_ is a random variable, and the probability with which extant species are sampled a parameter to estimate, which we denote by *f*_*p*_.

### Reconstructed tree

A realization of the diversification process from *t*_*c*_ to *t*_*p*_ can be represented graphically as a tree, see Figure 1. The complete tree shows all the species that have originated in the process (Fig. 1a). However, in practice we have only access to the reconstructed tree, i.e., the complete tree from which we remove all the species that became extinct before the present or that were not sampled (Fig. 1b). While it would be straight-forward to infer information about the diversification process based on the complete tree, this task is much more challenging when only the reconstructed tree is available.

**Fig. 1.**
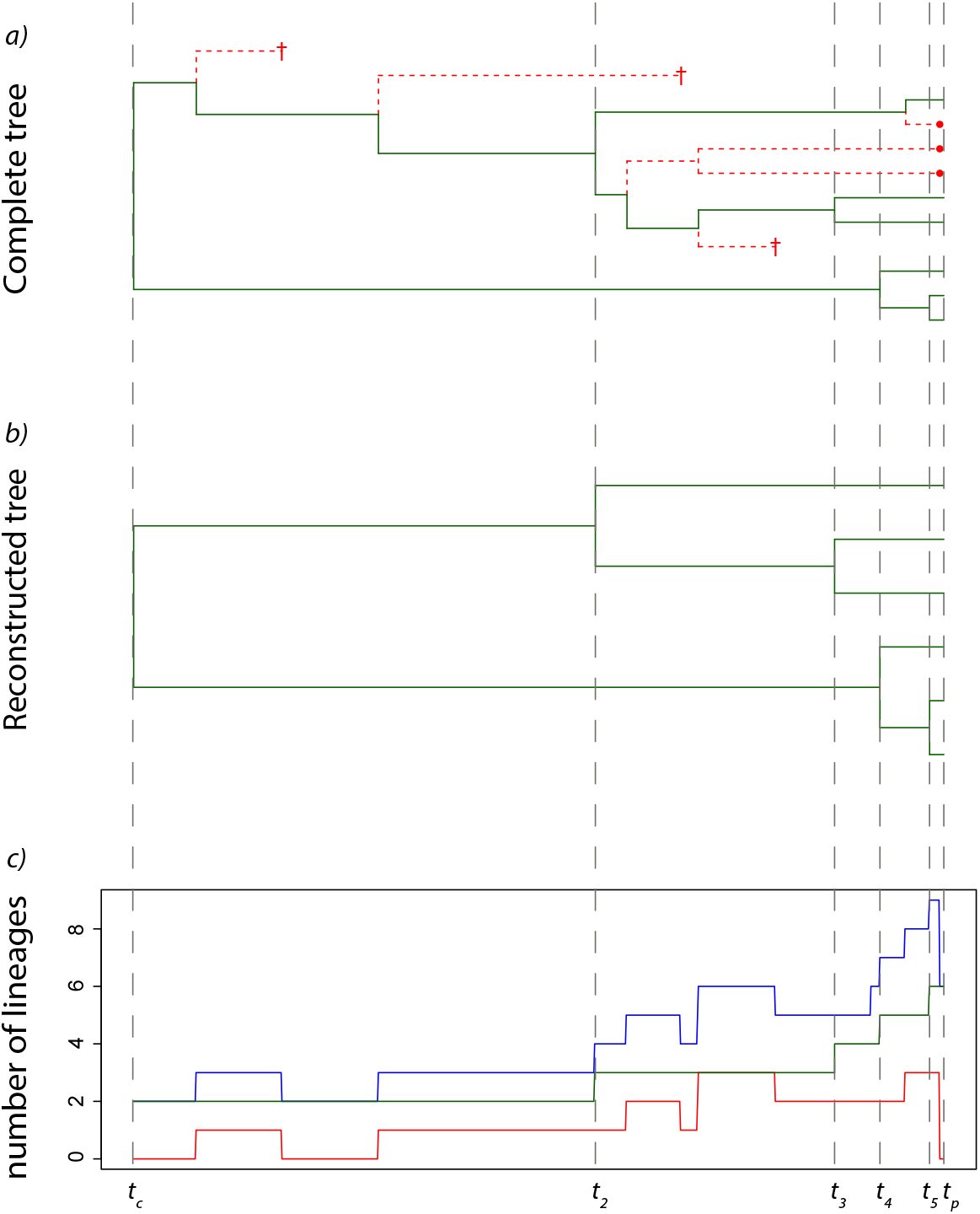
a) Full tree where missing species are plotted as red dashed lines: the ones ending in a cross become extinct before the present, whereas the ones ending with a red dot are unsampled species at the present; b) Corresponding reconstructed tree in which only extant species are present. This is the type of tree we usually work with because actual phylogenetic trees are usually obtained from molecular data taken from extant species; c) Lineages-through-time plot: the green line represents the number of lineages leading to extant species (*k*), the red line represents lineages leading to extinct or unsampled species (*m*), and the blue line represents the total number of lineages (*n* = *k* + *m*).

This paper deals with likelihood formulas for a reconstructed phylogenetic tree. The number of tips equals the number of sampled extant species *k*_*p*_. We assume that also the number of unsampled extant species is known, a number we denote by *m*_*p*_. The information contained in a phylogenetic tree consists of a topology and a set of branching times. For a large set of diversification models, including the diversity-dependent one, all trees having the same branching times but different topologies are equally probable [8]. Hence, we can discard the topology for the likelihood computation. We denote the vector of branching times by 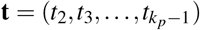, where *t*_*k*_ is the branching time at which the phylogenetic tree changes from *k* to *k* + 1 branches. It will be convenient to set *t*_0_ = *t*_1_ = *t*_*c*_ and 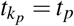.

### Likelihood conditioning

It is common practice to condition the likelihood on the survival of both ancestor lineages to the present time [9]. Indeed, we would only do an analysis on trees that have actually survived to the present. In particular, we impose the crown age of the reconstructed tree. This corresponds to conditioning on the event that both ancestor species have extant descendant species. We do not require that these descendant species have been sampled.

## 3 The framework of Etienne et al

Etienne et al. [3] presented an approach to compute the likelihood of a phylogeny for the diversity-dependent model. It is based on a new variable, 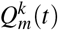, which they described as “the probability that a realization of the diversification process is consistent with the phylogeny up to time *t*, and has *n* = *m* + *k* species at time *t*” (Ref. [3], Box 1), where *k* lineages are represented in the phylogenetic tree (because they are ancestral to one of the *k*_*p*_ species extant and sampled at present) and *m* additional species are present but unobserved (Fig. 1c). These species might not be in the phylogenetic tree because they became extinct before the present or because they are either not discovered or not sampled yet (see introduction). From hereon we will refer to these species denoted by *m* as missing species. We cannot ignore these missing species, because in a diversity-dependent speciation process, they can influence the speciation and extinction rates.

We start by describing the computation of the variable 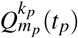, which proceeds from the crown age *t*_*c*_ to the present time *t*_*p*_. It is convenient to arrange the values 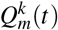, with *m* = 0, 1, 2,…, into the vector **Q**^*k*^(*t*). The initial vector 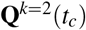 is transformed into the vector **Q**^*k*^(*t*) at a later time *t* as follows (Ref. [3], Appendix S1, Eq. (S1)):

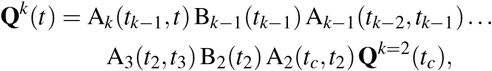

with *t*_*k*−1_ ≤ *t* ≤ *t*_*k*_. The operators *A*_*k*_ and *B*_*k*_ are infinite-dimensional matrices that operate along the tree, on branches and nodes, respectively (Fig. 2). Continuing this computation until the present time *t*_*p*_, we get

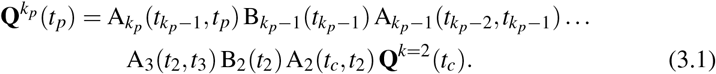

**Fig. 2.**
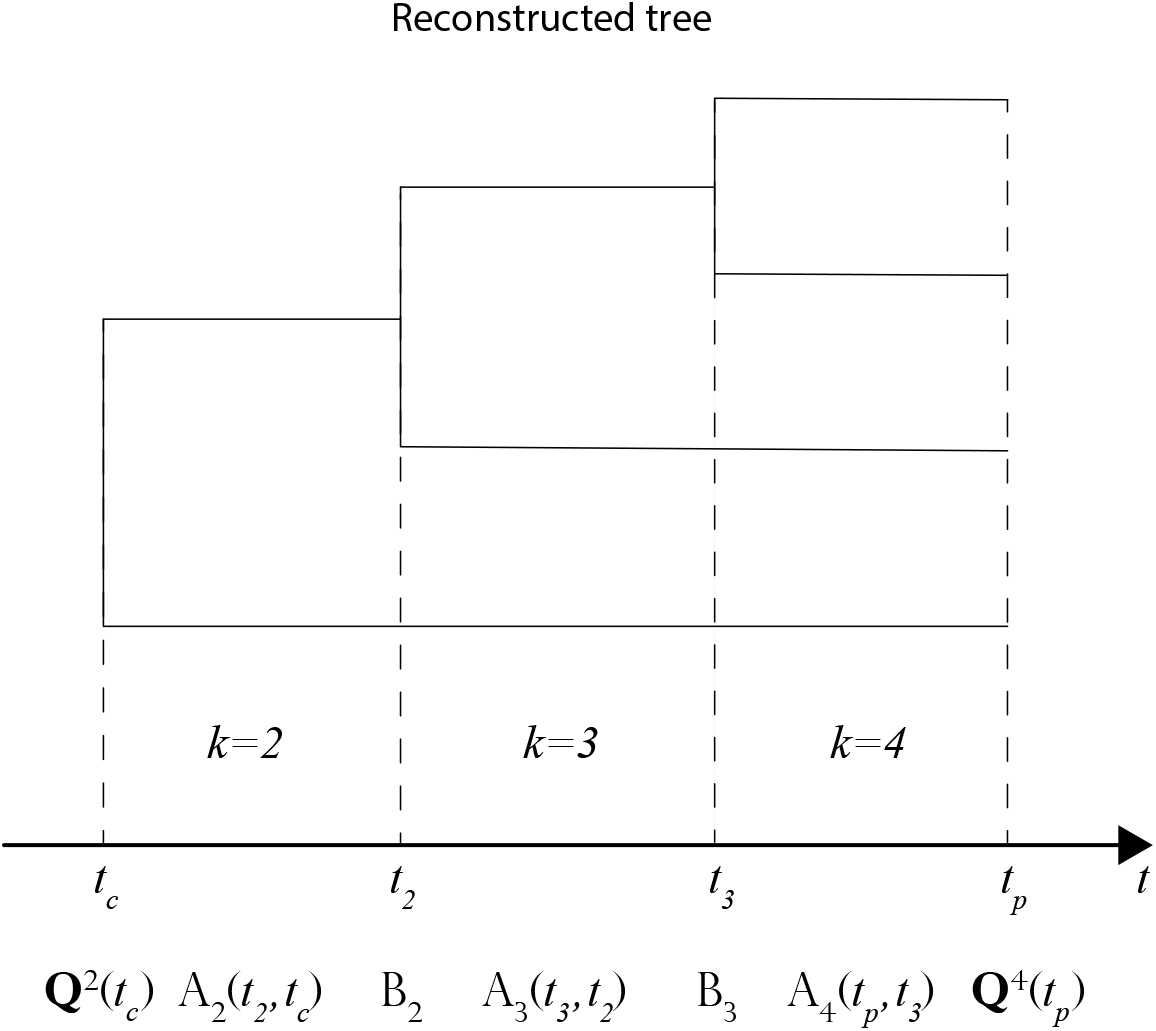
An example of how to build a likelihood for a tree with *k*_*p*_ = 4 tips. We start with a vector **Q**^2^(*t*_*c*_) at the crown age. We use A_*k*_(*t*_*k*_,*t*_k−1_) and B_*k*_(*t*_*k*_) to evolve the vector across the entire tree (on branches and nodes, respectively) up to the present time *t*_*p*_ according to **Q**^4^(*t*_4_) = *A*_4_(*t*_4_,*t*_3_)*B*_3_(*t*_3_)*A*_3_(*t*_3_,*t*_2_)*B*_2_(*t*_2_)*A*_2_(*t*_2_,*t*_*c*_)**Q**^2^(*t*_*c*_). At the present time the likelihood accounting for *m*_*p*_ missing species will be proportional to the *m*_*p*_-th component of the vector 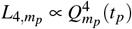.

Note that Eq. (3.1) generalizes Eq. (S1) of Ref. [3] to the case in which the rates are time-dependent.

We specify the different terms appearing in the right-hand side of Eq. (3.1):

– For the initial vector **Q**^*k*=2^(*t*_*c*_) we assume that there are no missing species at crown age, that is, 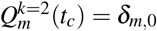.
– The matrix *A*_*k*_ corresponds to the dynamics of 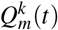 in the time interval [*t*_*k*−1_, *t*_*k*_], during which the phylogenetic tree has *k* branches. Etienne et al. [3] argued that these dynamics are given by the following ODE system (Ref. [3], Box 1, Eq. (B2)):

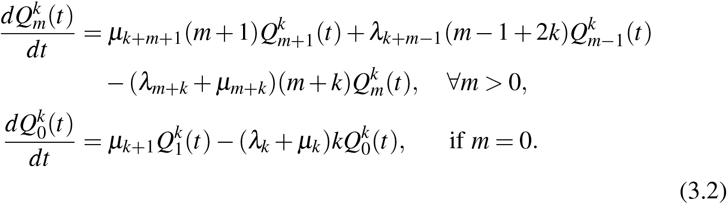 The quantity 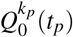 is the likelihood for the tree at the present time, assuming that all the species survived to the present have been sampled. We can collect the coefficients of 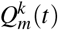 on the right-hand side of the ODE system in a matrix V_*k*_(*t*). If we do so, the system can be rewritten as

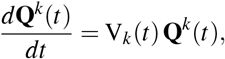

which has solution

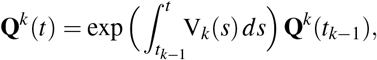

so that

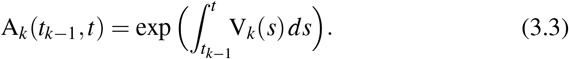
– The matrix B_*k*_ transforms the solution of the ODE system ending at *t*_*k*_ into the initial condition of the ODE system starting at *t*_*k*_. It is a diagonal matrix with components *kλ*_*k*+*m*_*dt*, so that

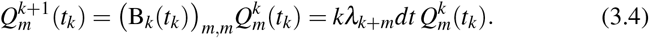 The multiplication by *λ*_*k*+*m*_*dt* corresponds to the probability that a speciation occurs in the time interval [*t*_*k*_, *t*_*k*_ + d*t*]. In the likelihood expressions we will omit the differential (a choice that is widely adopted across the vast majority of this kind of models in the literature) as it is actually not essential in parameter estimation. Therefore, we will work with a likelihood density, but for simplicity we will refer to it as a likelihood.

We are then ready to formulate the claim made by Etienne et al. [3] (in particular, see their Eqs. (S2) and (S6) in Appendix S1).

### Claim 31

*Consider the diversity-dependent diversification model, given by speciation rates λ*_*n*_(*t*) *and extinction rates μ*_*n*_(*t*). *The diversification process starts at crown age t*_*c*_ *with two ancestor species, and ends at the present time t*_*p*_, *at which a fixed number of species m*_*p*_ *are not sampled. A phylogenetic tree is constructed for the sampled species. Then, the likelihood that the phylogenetic tree has k*_*p*_ *tips and vector of branching times* 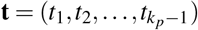, *conditional on the event that both crown lineages survive until the present, is equal to*

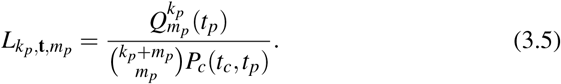

*The term* 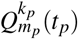 *in the numerator of this expression is obtained from Eq. (3.1)*. *The term P*_*c*_(*t*_*c*_, *t*_*p*_) *in the denominator, where the subscript c stands for conditioning, is equal to*

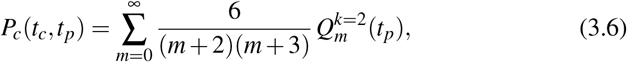

*where* 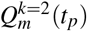 *is again obtained from Eq. (3.1)*.

The structure of the likelihood expression (3.5) can be understood intuitively. It is proportional to 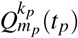, which in Etienne et al.’s interpretation is the probability that the diversification process generates the phylogenetic tree with *k*_*p*_ tips and *m*_*p*_ missing species at present time *t*_*p*_. The combinatorial factor 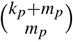 accounts for the number of ways to select *m*_*p*_ missing species out of a total pool of *k*_*p*_ + *m*_*p*_ species. The factor *P*_*c*_(*t*_*c*_, *t*_*p*_) is the probability that both ancestor species at crown age *t*_*c*_ have descendant species at the present time *t*_*p*_. Hence, this factor applies the likelihood conditioning.

Etienne et al. [3] provided numerical evidence that Claim 31 is in agreement with the likelihood provided by Nee et al. [9] under the hypothesis of diversity-independent speciation and extinction rates and no missing species at the present. However, a rigorous analytical proof, even for this specific case, has not yet been given. In this paper we show that Claim 31 holds (1) for the diversity-independent (but possibly time-dependent) case and (2) for the diversity-dependent case without extinction (i.e., extinction rate *μ* = 0).

## 4 The likelihood for the diversity-independent case

Claim 31 proposes a likelihood expression for the case with a known number of unsampled species at the present, i.e., it accounts for *n*-sampling. For the diversity-independent case, i.e., *λ*_*n*_(*t*) = *λ* (*t*) and *μ*_*n*_(*t*) = *μ*(*t*), the likelihood is contained in a more general result established by Lambert et al. [7]. In the following proposition we derive an explicit likelihood expression by restricting the result of Lambert et al. to the diversity-independent case.

### Proposition 1

*Consider the diversity-independent diversification model, given by speciation rates λ* (*t*) *and extinction rate μ*(*t*). *The diversification process starts at crown age t*_*c*_ *with two ancestor species, and ends at the present time t*_*p*_, *at which a fixed number of species m*_*p*_ *are not sampled. A phylogenetic tree is constructed for the sampled species. Then, the likelihood that the phylogenetic tree has k*_*p*_ *tips and vector of branching times* 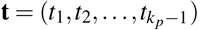, *conditional on the event that both crown lineages survive until the present, is equal to*

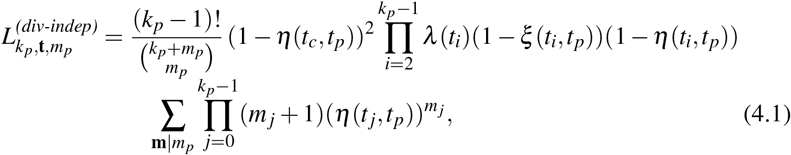

*where we used the convention t*_0_ = *t*_1_ = *t*_*c*_. *The components m*_*j*_ (*with j* = 0, 1,…, *k*_*p*_− 1*) of the vectors* **m**, *in the sum on the second line, are non-negative integers satisfying* 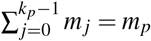. *The functions ξ* (*t*, *t*_*p*_) *and η*(*t*, *t*_*p*_) *are given by*

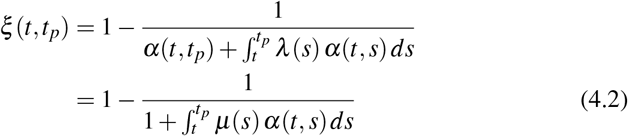

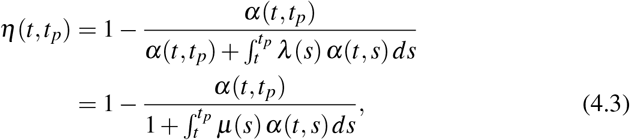

*with*

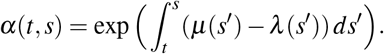

The functions *ξ* (*t*, *t*_*p*_) and> *η*(*t*, *t*_*p*_) are those appearing in Kendall’s solution of the birth-death model (see Ref. [6], Eqs. (10–12)), and are useful to describe the process when time-dependent rates are involved. Given the probability *P*_*n*_(*t*, *t*_*p*_) of realizing a process starting with 1 species at time *t* and ending with *n* species at time *t*_*p*_, we have that *ξ* (*t*, *t*_*p*_) = *P*_0_(*t*, *t*_*p*_) and 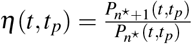 for any *n*^★^ > 0.

**Proof** The likelihood for *n*-sampling was originally provided by Ref. [7], Eq. (7), but we start from the explicit version provided in Ref. [4], Eq. (1), see corrigendum in Ref. [2],

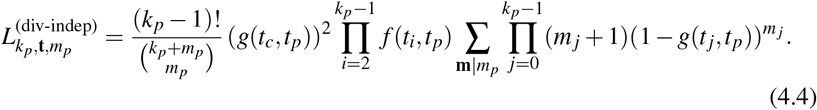

Lambert et al. [4, 7] specify the functions *f* (*t, t*_*p*_) and *g*(*t, t*_*p*_) as the solution of a system of ODEs for the case of protracted speciation, a model where speciation does not take place instantaneously but is initiated and needs time to complete. The standard diversification model is then obtained by taking the limit in which the speciation-completion rate tends to infinity. In this limit the four-dimensional system of Lambert et al. [4], Eq. (2), reduces to a two-dimensional system of ODEs,

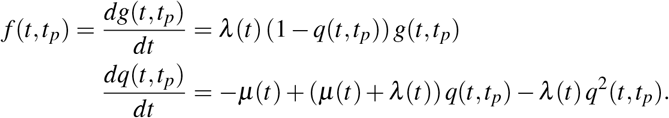

Note that in this paper time *t* runs from past to present rather than from present to past as in Lambert et al. [4]. The conditions at the present time *t*_*p*_ are given by *g*(*t*_*p*_, *t*_*p*_) = 1 and *q*(*t*_*p*_, *t*_*p*_) = 0.

The solution of this system of ODEs can be expressed in terms of *η*(*t, t*_*p*_) and *ξ* (*t, t*_*p*_),

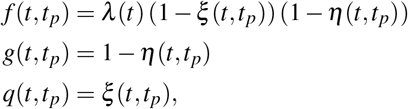

which can be checked using the derivatives of the expressions 4.3 and 4.2

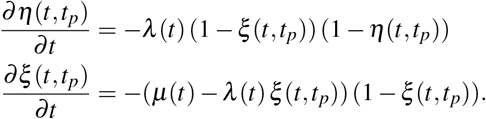

Substituting the functions *f* (*t, t*_*p*_) and *g*(*t, t*_*p*_) into the likelihood expression (4.4) concludes the proof.

The functions *ξ* (*t, t*_*p*_) and *η*(*t, t*_*p*_) are directly related to the functions used by Nee et al. [9]. In particular, the functions they denoted by *P*(*t, t*_*p*_) and *u_t_* correspond in our notation to 1 − *ξ* (*t, t*_*p*_) and *η*(*t, t*_*p*_), respectively.

This correspondence allows us to get an intuitive understanding of the likelihood expression (4.1). First consider the case without missing species. Setting *m*_*p*_ = 0, we get

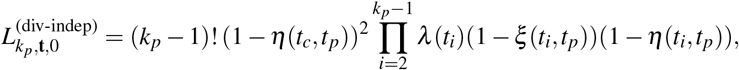

which is identical to the breaking-the-tree likelihood of Nee et al. (Ref. [9], Eq. (20)). In the latter approach the phylogenetic tree is broken into single branches: two for the interval [*t*_*c*_, *t*_*p*_] and one for each interval [*t*_*i*_, *t*_*p*_] with *i* = 2, 3, …, *k*_*p*_ − 1. Each branch contributes a factor (1 −*ξ* (*t*_*i*_, *t*_*p*_))(1 −*η*(*t*_*i*_, *t*_*p*_)), equal to the probability that the branch starting at *t*_*i*_ has a single descendant species at *t*_*p*_. For the two branches originating at *t*_*c*_, the factor (1 −*ξ* (*t*_*i*_, *t*_*p*_)), equal to the probability of having (one or more) descendant species, drops due to the conditioning. For the other branches, there is an additional factor *λ* (*t*_*i*_) for the speciation events.

Next consider the case with missing species. Each of the branches resulting from breaking the tree can contribute species to the pool of *m*_*p*_ missing species. For the branch over the interval [*t*_*j*_, *t*_*p*_], there are *m*_*j*_ such species, each contributing a factor *η*(*t*_*j*_, *t*_*p*_) to the likelihood. Indeed, 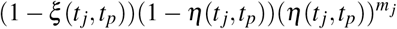 is equal to probability of having exactly *m*_*j*_ + 1 descendant species at the present time. One of these species is represented in the phylogenetic tree, justifying the combinatorial factor (*m*_*j*_ + 1) in the second line of Eq. (4.1).

Finally, we recall the expressions for the functions *ξ* (*t, t*_*p*_) and *η*(*t, t*_*p*_) in the case of constant rates, *λ* (*t*) = *λ* and *μ*(*t*) = *μ*,

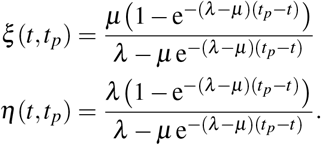

## 5 Equivalence for the diversity-independent case

Likelihood formula (4.1) allows speciation and extinction rates to be arbitrary functions of time, *λ* (*t*) and *μ*(*t*). Here we show that, for the diversity-independent case, we find the same likelihood formula with the approach of Etienne et al. [3].

### Theorem 1

*Claim 31 holds for the diversity-independent case*

**Proof** The proof relies heavily on generating functions. First, we introduce the generating function for the variables 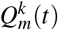,

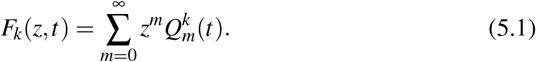

The set of ODEs satisfied by 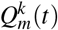, Eq. (3.2), transforms into a partial differential equation (PDE) for the generating function *F*_*k*_(*z, t*),

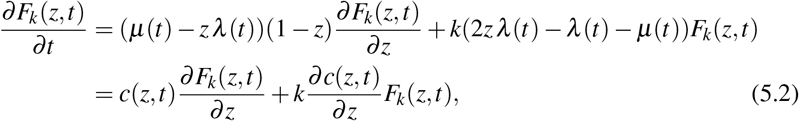

with

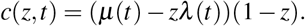

Note that the number of branches *k* changes at each branching time, so that the PDE for *F*_*k*_(*z, t*) is valid only for *t*_*k*−1_ ≤ *t* ≤ *t*_*k*_ (corresponding to the operator *A*_*k*_). At branching time *t*_*k*_, the solution *F*_*k*_(*z, t*_*k*_) has to be transformed to provide the initial condition for the PDE for *F*_*k*+1_(*z, t*) at time *t*_*k*_ (corresponding to the operator *B*_*k*_). Using Eq. (3.4) and dropping the differential, we get

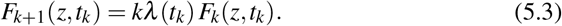

The initial condition at crown age is *F*_2_(*z, t_c_*) = 1 because 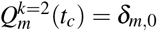.

Next, we define *P*_*n*_(*s, t*) as the probability that the birth-death process that started with one species at time *s* has *n* species at time *t*. The corresponding generating function is defined as,

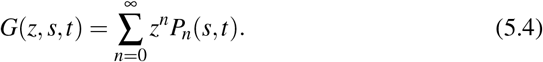

The set of ODEs satisfied by *P*_*n*_(*s, t*), Eq. (2.1), transforms into a PDE,

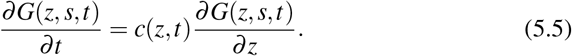

Its solution was given by Kendall (Ref. [6], Eq. (9)),

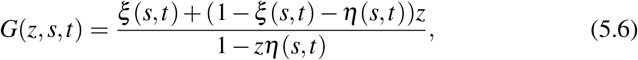

where *ξ* (*s, t*) and *η*(*s, t*) are given in Eqs. (4.2) and (4.3).

The generating function *F*_*k*_(*z, t*) can be expressed in terms of the generating function *G*(*z, s, t*), as shown in the following lemma.

### Lemma 1

*The generating function F*_*k*_(*z, t*) *of the variables* 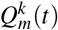 *is given by*

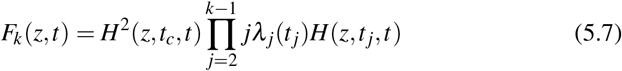

*with*

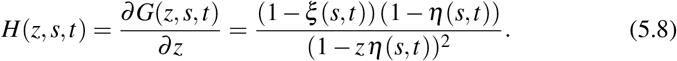

To prove the lemma, let us suppose that the solution of Eq. (5.2) is of the form,

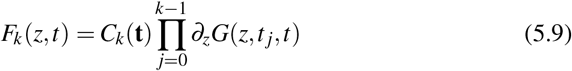

where *C*_*k*_(**t**) is a constant depending on the branching times. We used the convention *t*_0_ = *t*_1_ = *t*_*c*_ and the short-hand notation *∂*_*z*_ for the partial derivative with respect to *z*. This expression can be rewritten as,

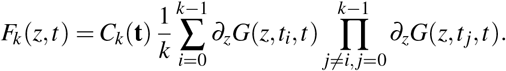

The partial derivatives of *F*_*k*_ can now be computed,

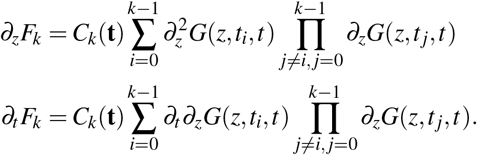

We substitute these expressions into the PDE, Eq. (5.2),

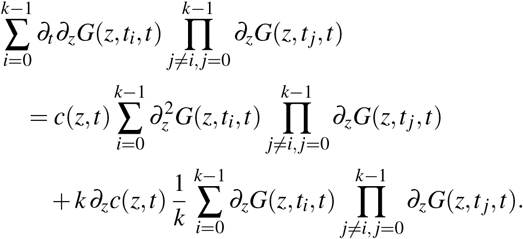

This equation is satisfied if the following equation is satisfied for every *i* = 0, 1,…, *k*,

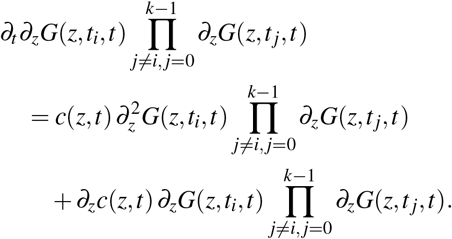

This is the case if

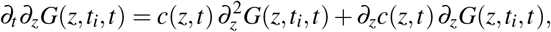

or, equivalently, if

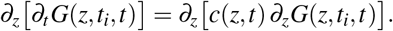

This is an identity because *G*(*z, t*_*i*_, *t*) satisfies Eq. (5.5).

Next, we verify that the constants *C*_*k*_(**t**) can be determined such that initial conditions (5.3) are satisfied. This is indeed the case if we take

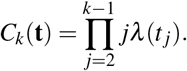

Introducing the function *H*(*z, s, t*) and using *t*_0_ = *t*_1_ = *t*_*c*_ complete the proof of the lemma.

Next, we use Eq. (5.7) to derive an explicit expression for the likelihood (3.5) of Claim 31. It will be useful to have explicit expressions for derivatives of the function *H*(*z, s, t*). It follows from Eq. (5.8) that

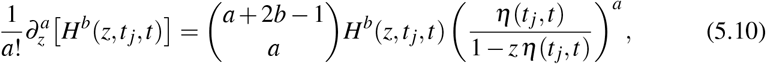

where *a* and *b* are positive integers.

To evaluate the numerator of Eq. (3.5), we have to extract 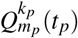 from the generating function 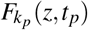. Using Leibniz’ rule,

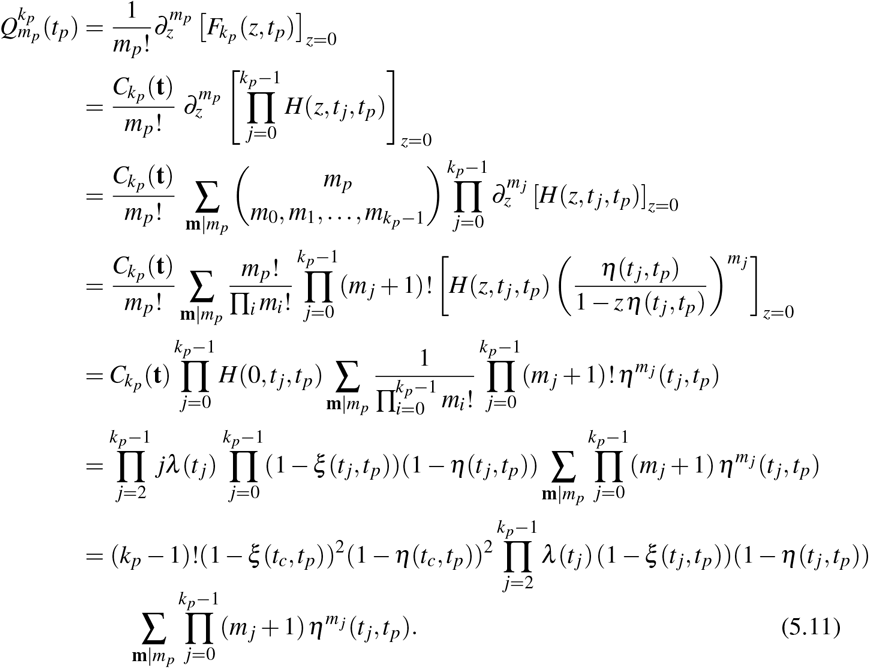

To evaluate the denominator of Eq. (3.5), we have to extract 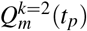 from the generating function,

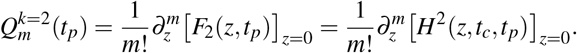

Substituting into Eq. (3.6) and using Eq. (5.10), we get

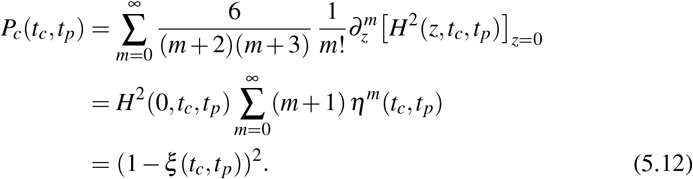

Finally, substituting Eqs. (5.11) and (5.12) into the likelihood formula (3.5) of Claim 31,

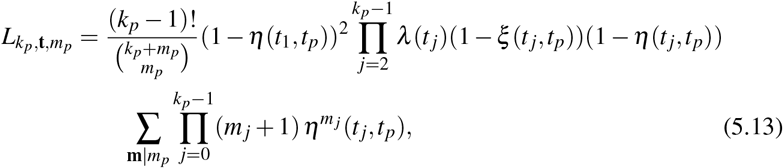

which is identical to likelihood formula (4.1). This concludes the proof of the theorem.

## 6 A note on sampling a fraction of extant species

Nee et al. [9] noted that one way to model the sampling of extant species is equivalent to a mass extinction just before the present. This sampling model corresponds to sampling each extanct species with a given probability *f*_*p*_, which has also been called *ρ*-sampling [7]. We use the link with mass extinction to extend the previous formula for *n*-sampling to the case of *ρ*-sampling.

First, we formulate the *ρ*-sampling version of Claim 31.

### Claim 61

*Consider the diversity-dependent diversification model, given by speciation rates λ_n_*(*t*) *and extinction rates μ_n_*(*t*). *The diversification process starts at crown age t*_*c*_ *with two ancestor species, and ends at the present time t*_*p*_, *at which extant species are sampled with probability f*_*p*_. *Then, the likelihood of a phylogenetic tree with k*_*p*_ *tips and branching times* **t**, *conditional on the event that both crown lineages survive until the present, is equal to*

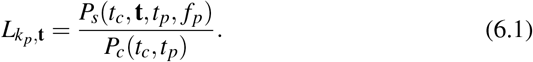

*The term P*_*s*_(*t*_*c*_, ***t***, *t*_*p*_, *f*_*p*_) *in the numerator, where the subscript s stands for sampling, is equal to*

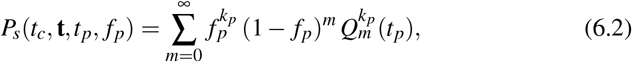

*where* 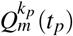 *is obtained from Eq. (3.1)*. *The term P*_*c*_(*t*_*c*_, *t*_*p*_) *in the denominator, where the subscript c stands for conditioning, is equal to*

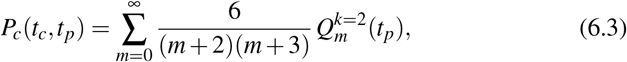

*where* 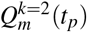 *is again obtained from Eq. (3.1)*.

Next, we establish as a reference the likelihood formula for *ρ*-sampling in the diversity-independent case.

### Proposition 2

*Consider the diversity-independent diversification model, given by speciation rates λ* (*t*) *and extinction rates μ*(*t*). *The diversification process starts at crown age t*_*c*_ *with two ancestor species, and ends at the present time t*_*p*_, *at which extant species are sampled with probability f*_*p*_. *Then, the likelihood of a phylogenetic tree with k*_*p*_ *tips and branching times* **t**, *conditional on the event that both crown lineages survive until the present, is equal to*

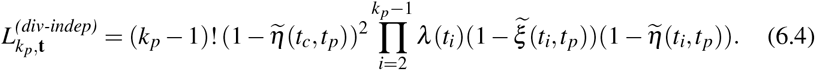

*The functions* 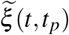 *and* 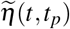 *are given by*

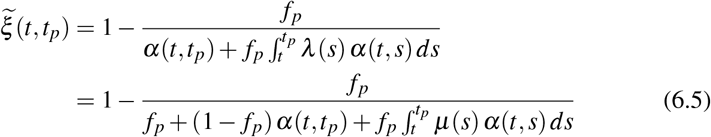

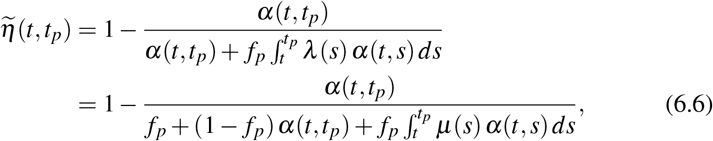

*with*

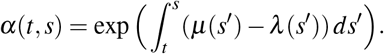

**Proof** We use the equivalence between *ρ*-sampling and a mass extinction, see Ref. [9], Eq. (31). We introduce a modified extinction rate *μ*(*t*) containing a delta function just before the present,

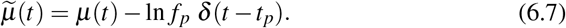

The likelihood formula is then obtained by setting *m*_*p*_ = 0 in Eq. (4.1), while evaluating the functions *ξ* (*t, t*_*p*_) and *η*(*t, t*_*p*_) with the modified extinction rate 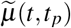. This establishes Eq. (6.4); it remains to be proven that the modified functions 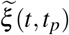 and 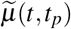 are given by Eqs. (6.5) and (6.6). This follows by noting that the modified version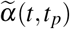 of the function *α*(*t, t*_*p*_) appearing in Eqs. (4.2) and (4.3) satisfies

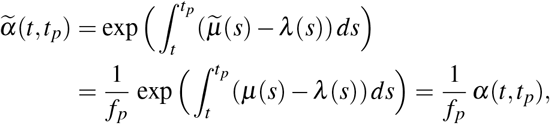

while 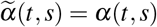 if *s* < *t*_*p*_.

We are then ready to establish the following result.

### Theorem 2

*Claim 61 holds for the diversity-independent case*

**Proof** We use again the equivalence between *ρ*-sampling and a mass extinction, see Eq. (6.7). Due to Theorem 1, likelihood formula (3.5) is valid for the diversity-independent case. Hence, we can derive the corresponding likelihood formula for *ρ*-sampling by introducing the modified extinction rate 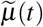, and setting *m*_*p*_ = 0 in the likelihood formula for *n*-sampling.

The introduction of the modified extinction rate 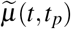 corresponds to applying an additional operator to the vector 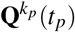 at the present time. In particular, the modified vector 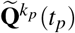 is given by

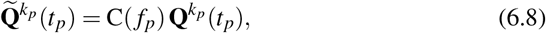

where the operator C(*f*_*p*_) corresponds to the following ODE, acting in a small time interval [*t*_*p*_ − *ε*, *t*_*p*_] before the present,

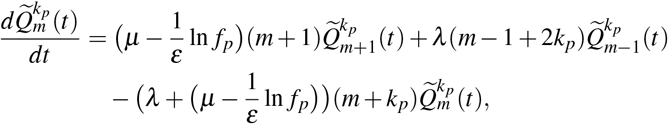

where we added a delta peak to the extinction rate, Eq. (6.7), in the ODE satisfied by **Q**^*k*^(*t*), Eq. (3.2). In the limit *ε* → 0 the terms in 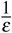 dominate, so that

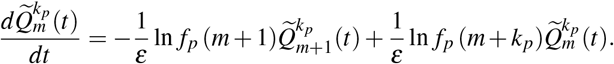

This can be rewritten in matrix form as

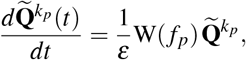

where the operator W(*f*_*p*_) is an infinite-dimensional matrix with components

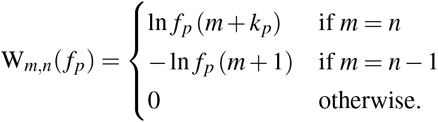

Hence, the operator C(*f*_*p*_), which is also an infinite-dimensional matrix, is equal to

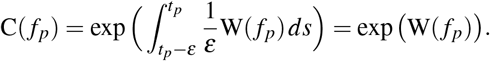

We need the row *m* = 0 to evaluate the likelihood, which is equal to

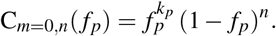

We are then ready to evaluate likelihood formula (3.5) with the modified extinction rate. Setting *m*_*p*_ = 0, we get

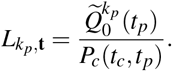

Recall that the conditioning probability *P*_*c*_(*t*_*c*_, *t*_*p*_) is not affected by the process of sampling extant species. We get

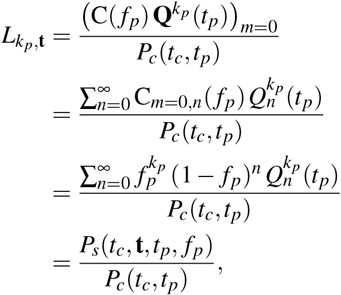

which is identical to Eq. (6.4). This ends the proof.

Finally, we give the expressions for the functions 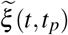 and 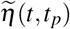 in the case of constant rates, *λ* (*t*) = *λ* and *μ*(*t*) = *μ*,

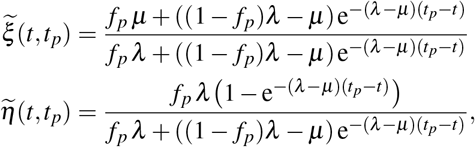

which are identical to Eqs. (4) and (5) in the paper by Stadler [11].

## 7 The diversity-dependent case without extinction

Rabosky & Lovette [10] derived the likelihood for a particular instance of the diversity-dependent diversification model, namely, when there is no extinction. This is the only case for which a diversity-dependent likelihood formula is available. Here we show that this case is dealt with correctly in the approach of Etienne et al. [3].

We start by reformulating the result of Rabosky & Lovette [10] in our notation.

### Proposition 3

*Consider the diversity-dependent model without extinction, given by speciation rates λ*_*n*_(*t*). *The diversification process starts at crown age t*_*c*_ *w ith two ancestor species, and ends at the present time t*_*p*_, *at which all extant species are sampled. Then, the likelihood of a phylogenetic tree with k*_*p*_ *tips and branching times* **t** *is equal to*

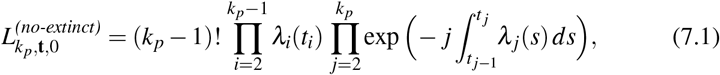

*where we used the convention t*_1_ = *t*_*c*_ *and* 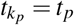.

**Proof** Eq. (7.1) follows from Eqs. (2.4) and (2.5) in Ref. [10], by noting that *ξ*_*i*_ in their notation corresponds to

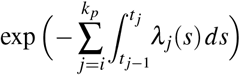

in our notation.

Note that in the case without extinction likelihood conditioning has no effect.

### Theorem 3

*Claim 31 holds for the diversity-dependent case without extinction*

**Proof** To evaluate likelihood expression (3.5), we have to solve the ODE for 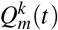, Eq. (3.2). Because species cannot become extinct and because all extant species are sampled, every species created during the process is represented in the phylogeny, i.e., there are no missing species. Hence, only the *m* = 0 component of **Q**^*k*^(*t*) is different from zero. The ODE simplifies to where

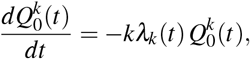

*t* belongs to [*t*_*k*−1_, *t*_*k*_]. Note that in this time interval there are exactly *k* species. Given the initial condition 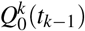 at *t*_*k*−1_, the solution is

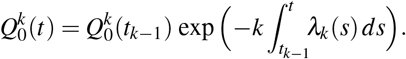

At branching time *t*_*k*_, variable 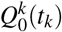 is transformed into variable 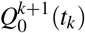,

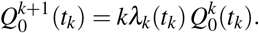

Using the initial condition at crown age *t*_*c*_, 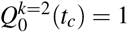, we get

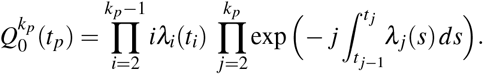

Substituting into Eq. (3.5) yields the desired result.

## 8 Concluding remarks

We have shown here that for the diversity-independent, but time-dependent birth-death model with *n*-sampling, the framework of Etienne et al. [3] yields the same likelihood as derived by Lambert et al. ([4],[2]). This provides strong support for the correctness of this framework, but does not prove that it is also correct for the case of diversity-dependence. We have thus far not been able to provide alternative evidence for this framework, apart from the fact that parameter estimations on simulations of this model provide reasonable, although sometimes biased, estimates [3]. We hope that our analysis here will suggest directions for a further substantiating of the framework.

Most existing macroevolutionary models rely on the hypothesis that the subcomponents of trees do not interact (and one can thus apply a breaking-the-tree approach, as in [9], pag. 308), therefore letting the likelihood be a factorization of terms that comes independently from the tree’s edges and nodes. However, such a hypothesis is not always valid. The diversification process likely also depends on properties of other lineages than the lineage under consideration. The analytical treatment of Etienne et al.’s [3] arguments presented in this work suggests a direction towards deriving the likelihood for much more complicated models with “interacting branches”, with the arguably simplest case being diversity dependence, i.e. dependence only on the total number of lineages present at any time. Our work, showing analytically that Etienne et al.’s model agrees with existing formulas for likelihoods of simple diversification models, suggests that future models that aim to deal with interacting branches should consider such a structure as a reference point, in the same fashion as models dealing with “breakable” trees often refer to Nee et al.’s [9] paradigm.

In this article we have proved that the framework to compute a likelihood for diversity-dependent processes by Etienne et al. [3] agrees with analytical results obtained for diversity-independent diversification models. This suggests that the framework is valid for more general models that take into account the effect of diversity of speciation and extinction rates while still being able to deal with unsampled species in the phylogeny, when this number is known. Our results can thus improve the understanding of the general architecture of macroevolutionary diversification models providing useful tools for the development of new models.

## References

1. Bailey, N.T.: The elements of stochastic processes with applications to the natural sciences. John Wiley & Sons(1990)

2. Etienne, R.S.: Corrigendum. Evolution (2017). DOI:10.1111/evo.13314

3. Etienne, R.S., Haegeman, B., Stadler, T., Aze, T., Pearson, P.N., Purvis, A., Phillimore, A.B.: Diversity-dependence brings molecular phylogenies closer toagreement with the fossil record. Proc. R. Soc. Lond. B: Biol. Sci. 279(1732), 1300–1309 (2012)

4. Etienne, R.S., Morlon, H., Lambert, A.: Estimating the duration of speciation from phylogenies. Evolution 68(8), 2430–2440 (2014)

5. Etienne, R.S., Rosindell, J.: Prolonging the past counteracts the pull of the present: protracted speciation can explain observed slowdowns in diversification. Syst. Biol. 61(2), 204–213 (2011)

6. Kendall, D.G.: On the generalized “birth-and-death” process. Ann. Math. Stat. 19(1), 1–15 (1948)

7. Lambert, A., Morlon, H., Etienne, R.S.: The reconstructed tree in the lineage based model of protracted speciation. J. Math. Biol. 70(1/2), 367–397 (2015)

8. Lambert, A., Stadler, T.: Birth–death models and coalescent point processes: The shape and probability of reconstructed phylogenies. Theor. Pop. Biol. 90, 113–128 (2013)

9. Nee S.,May R. M. & Harvey P. H.: The reconstructed evolutionary process. Phil. Trans. R. Soc. Lond.B 344(1309), 305–311 (1994)

10. Rabosky, D.L., Lovette, I.J.: Density-dependent diversification in North American wood warblers. Proc. R. Soc. Lond. B: Biol. Sci. 275(1649), 2363–2371 (2008)

11. Stadler, T.: How can we improve accuracy of macroevolutionary rate estimates? Syst. Biol. 62(2), 321–329 (2013)

